# Development and validation of five native Aotearoa New Zealand genera-specific large brown macroalgal eDNA ddPCR assays

**DOI:** 10.1101/2025.03.21.644472

**Authors:** Mari Deinhart, Amber Brooks, Lisa Smith, Roberta D’Archino, Robert Hickson, Giuseppe C. Zuccarello, Scott D. Nodder, Jaret Bilewitch

## Abstract

Environmental DNA is a rapidly growing methodology for ecological studies, leading to an increase in innovative techniques, such as the droplet-digital PCR (ddPCR), that can quantify and target DNA at a high rate of sensitivity for eDNA studies. Large brown macroalgae, Phaeophyceae, are an ecologically significant, diverse, and abundant benthic group in temperate-reef coastal environments. Members of Phaeophyceae, such as Laminariales and Fucales, are becoming recognized as potential significant contributors to Blue Carbon, although measurement of their contribution is challenging. The goal of the present study was to track the fate of specific large, brown macroalgae (kelp) genera detritus, by designing and validating five assays targeting the eDNA of kelp genera that are abundant in the coastal environments of Aotearoa New Zealand: *Carpophyllum, Cystophora, Ecklonia, Lessonia*, and *Macrocystis*. Short, diagnostic sequences of the mitochondrial cytochrome c oxidase subunit III gene (*cox*3) were targeted using ddPCR analysis. Specificity and sensitivity testing were conducted to validate the design of each assay for future use. All five assays produced no cross-amplification and had limits of detection (LoD) that were less than 1 copy/µl. These novel, highly sensitive assays will be beneficial for future experimental and *in situ* eDNA studies that focus on any of these indigenous genera.

## 1. Introduction

Characterization of eDNA is a growing field for ecological and monitoring projects because sampling is non-invasive to organisms (Budd et al., 2021; Budd et al., 2023) and can be used for several types of objectives (Larson et al., 2020; Suren et al., 2024). There are several molecular methods to conduct eDNA surveys for projects that focus on ecological monitoring, biodiversity studies, conservation efforts, marine carbon dioxide removal (mCDR), such as coastal Blue Carbon (BC) strategies (Anglès d’Auriac et al., 2021; Frigstad et al., 2021; Monuki et al., 2021; Wesselmann et al., 2022), and species distributions (Bird et al., 2024; Budd et al., 2021; Budd et al., 2023; Deiner et al., 2017; Evans et al., 2016; Everts et al., 2021; McClenaghan et al., 2020; Monuki et al., 2021; Norgaard et al., 2021; Oladi et al., 2022; Pratomo et al., 2022; Sanchez et al., 2022; Zainal Abidin et al., 2022). A method that is being applied more recently to eDNA surveys is droplet digital PCR (ddPCR). The ddPCR method gives accurate and sensitive detection of eDNA present in a sample (Cao et al., 2020; Dimond et al., 2022; Everts et al., 2021; Miotke et al., 2014; Uchiyama et al., 2016; Wood et al., 2019), compared to real-time PCR (qPCR) and general eDNA metabarcoding (Wood et al., 2019). Along with those capabilities, ddPCR can quantify target DNA in a sample (Cao et al., 2020; Dimond et al., 2022; Everts et al., 2021; Miotke et al., 2014; Uchiyama et al., 2016; Wood et al., 2019), however this specific study will not be addressing this topic. These abilities make ddPCR a promising tool for studies aiming to quantify how much material may be present in the environment of its target DNA (Long & Berkemeier, 2020).

Phaeophyceae (Chromista), or brown macroalgae, are a diverse and unique group of organisms that provide numerous ecological services to marine environments around the globe. Members of Laminariales are classified as true kelp and are predicted to cover 1.71 million km^2^ globally, with currently, 138 accepted species (Fragkopoulou et al., 2022; Guiry, 2024). Fucales hold significant ecological roles and are estimated to have an aerial coverage of 2.57 million km^2^ globally (Fragkopoulou et al., 2022), with over 550 accepted species (Bringloe et al., 2020; Guiry, 2024). Members of the order Fucales are not considered ‘true’ kelp by some phycologists but other phycologists do classify them as kelp. (Fraser, 2012). Therefore, members of both orders will be referred to collectively as ‘kelp’ hereafter in this study. Together, the Laminariales and Fucales cover 25% of global coasts and are recognized as environmental engineers that drive the dynamics of their coastal marine environments (Starko et al., 2019; Steinberg et al., 1995; Steneck et al., 2002). In addition, they are regarded as marine foundational species due to their dominance in large bioregions, fast growth rates, high productivity, and habitat provision for numerous endemic and iconic marine species (Pessarrodona, Assis, et al., 2022; Pessarrodona, Filbee-Dexter, et al., 2022; Thomsen et al., 2024). More recently, it has been hypothesized that kelp are significant contributors to deep-sea carbon sequestration, once dislodged from their rocky reef habitats (Froehlich et al., 2019).

Aotearoa New Zealand (NZ) has a large marine 200 nautical mile exclusive economic zone (EEZ) covering approximately 4.3 million km^2^, including rich marine biodiversity (Costello et al., 2010; Gordon et al., 2010). NZ is the type locality for an estimated 164 Phaeophyceae species in 82 genera, with 32 recognized species of Fucales and 10 recognized Laminariales species (Nelson et al., 2023) (Preuss & Zuccarello, 2024), which (Schiel, 1990) provide habitat for a multitude of other native and endemic species (Hurd et al., 2004; Preuss & Zuccarello, 2024). The abundance of NZ kelp populations are changing at an accelerating pace (Blain et al., 2021; Mason & Blain, 2024; Montie et al., 2023; Tait et al., 2021; Thoral et al., 2023). Therefore, eDNA-based monitoring can also be a valuable tool for researching the changes in kelp populations and its role in BC (Gordon et al., 2010; Law et al., 2017; Spyksma et al., 2024; Tait et al., 2021).

The aim of this study was to develop ddPCR primer assays for eDNA of five abundant indigenous NZ kelp genera, with a focus on *Carpophyllum maschalocarpum* (Fucales), *Cystophora torulosa* (Fucales), *Ecklonia radiata* (Laminariales), *Lessonia variegata* (Laminariales), and *Macrocystis pyrifera* (Laminariales). These five genera can be found in rocky intertidal and subtidal habitats, fulfilling environmental services in different geographical locations in NZ. *Carpophyllum* is an endemic genus to NZ with six accepted species (Guiry, 2024) and is an abundant genus in NZ’s subtidal coastal zones (Dromgoole, 1973; Glombitza & Li, 1991a, 1991b; Neill & Nelson, 2016; Zhang et al., 2020). *Cystophora* has four species present in NZ, including *C. torulosa* which is found throughout the NZ upper subtidal to subtidal zone, as well as exposed coast*s*, (Neill & Nelson, 2016) (Pessarrodona & Grimaldi, 2022). *Ecklonia* has one species, *E. radiata*, occurring in NZ. *Lessonia* and *M. pyrifera* are found around NZ’s marine coastal environments creating unique temperate reefs that have historically provided valued ecosystem services to both Māori and European settlers (Hurd et al., 2004; Neill & Nelson, 2016; Nelson, 2020). *Lessonia* has seven accepted species in NZ, but only *L. variegata* occurs in Pōneke Wellington and Te Moana-o-Raukawa Cook Strait (Zuccarello & D’Archino, 2024).

The five assays developed in this study will be used for studies by the authors to assist in quantifying the contribution of each genus to BC in marine sediments and seawater in NZ. The development of these assays included optimisation and validation to identify the specific target genera, as well as to quantify the abundance of the targeted species in sediment and seawater samples. Although a designated species was used for each target assay design, the assays, excluding *M. pyrifera*, were not tested for cross-amplification of congeners. Consequently, the assays presented in this study are referred-to as ‘genera-specific’.

## 2. Materials & Methods

### 2.1. Sequence library & assay designs

Samples of 11 Phaeophyceae species were collected throughout the Te Whanga603nui-a-Tara Wellington region (Table 1) and were extracted using NucleoSpin® Plant II Kits (Macherey-Nagel, Germany), following the manufacturer’s protocol. Five of the eleven species represented the intended target species with six species of other brown macroalgae, distributed throughout the Te Whanganui-a-Tara Wellington coastline, were included as references and for specificity testing. These species were chosen due to their abundance in the region and ecological prominence. Extracted DNA was quantified using a quant-iT™ PicoGreen® dsDNA Assay kit (Molecular Probes Inc., Eugene, USA). DNA extracts were normalized to 1ng/µl and then PCR-amplified with primers for the mitochondrial cytochrome c oxidase subunit III protein coding gene (*cox*3), *cox*3-44F (CATCGCCACCCATTTCAT) & *cox*3-70039R (CATCGACAAAATGCCAATACCA) (Silberfeld et al., 2010). The PCR amplification profile for the eleven species followed Silberfeld et al. (79). Successfully amplified samples were submitted for bidirectional Sanger DNA sequencing (Macrogen Inc., Republic of Korea) using the same primers (Suppl. Table 1).

**Table 1.**
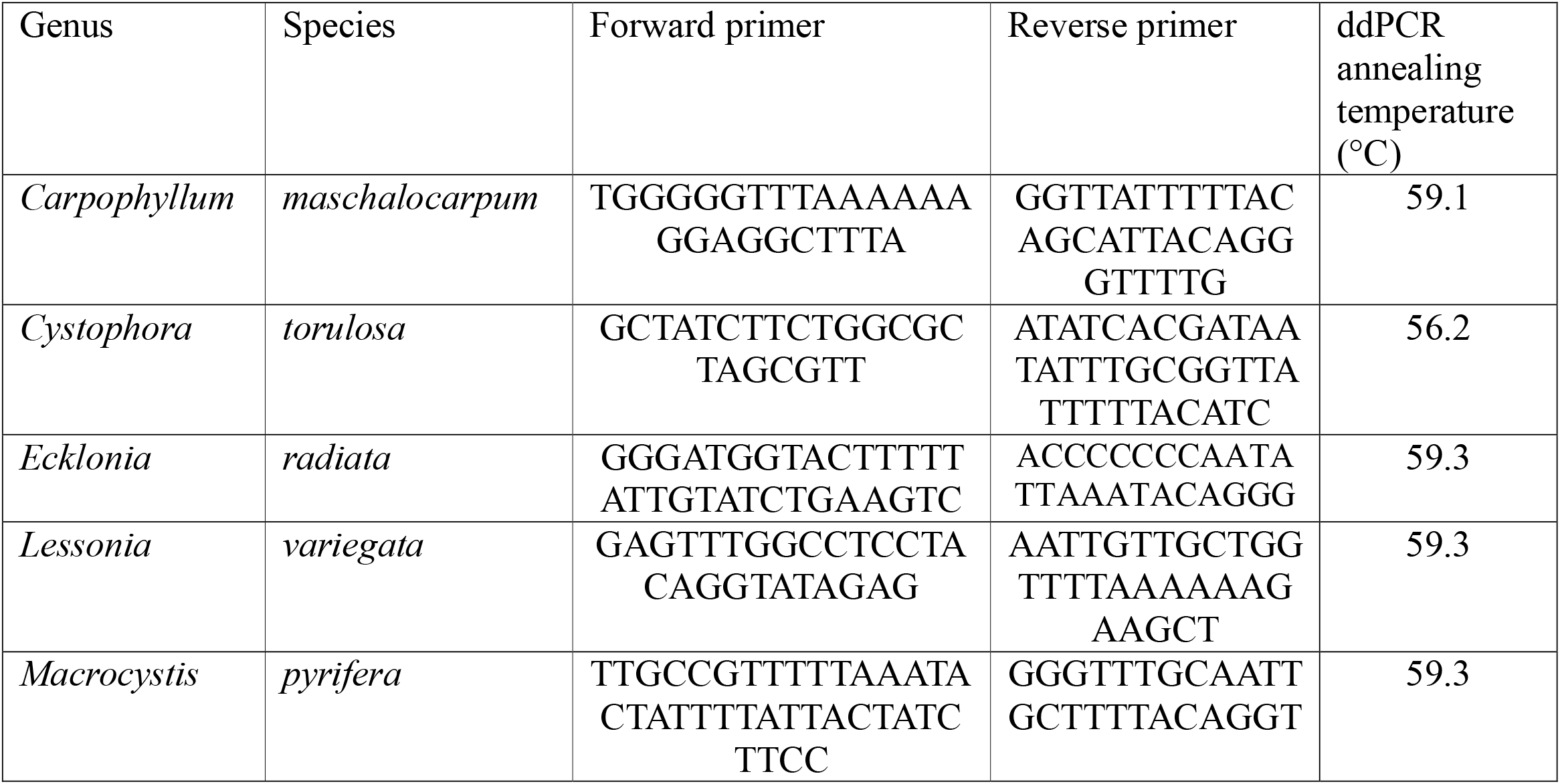
Summary of each genus primer assay design in this study. The target genus, its forward and reverse primer sequences (5’-3’), the optimal annealing temperature.

Forward and reverse sequences were assembled using Geneious Prime v11.0.9+11 (https://www.geneious.com)(Kearse et al., 2012). *Blastn* searches were used to confirm the identity of *cox*3 nucleotide sequences and DNA alignments against selected reference sequences in the National Center of Biotechnology Information database (NCBI; https://www.ncbi.nlm.nih.gov/nucleotide/) (Altschul et al., 1990). Sequence alignments were created using the MUSCLE plugin (Edgar, 2004) in Geneious Prime.

Primers for SYBR- and EvaGreen-based qPCR and ddPCR assays (respectively) for each of the five targeted genera were designed based on diagnostic variable regions (autapomorphies) of the *cox*3 alignment in Geneious. Published sequences of conspecifics, congeners and other target genera were obtained from NCBI and were aligned to identify unique short (<200bp) sequences for the five target species in this study (Suppl. Table 1).

### 2.2. Test conditions-qPCR

Primers were initially tested and optimized on a SYBR-based qPCR platform (Bio-Rad CFX) prior to re-optimization and further validation using EvaGreen dye with the Bio-Rad QX200 ddPCR system. Reactions contained 10 µl SsoAdvanced Universal SYBR Green Supermix (Bio-Rad Laboratories), 0.5µM of each primer, and 1 µl of DNA extract in a 20 µl total reaction volume. Each DNA sample was normalized to 1 ng/µl prior to analyses. Amplification was performed using a BioRad CFX qPCR system (Bio-Rad Laboratories), with data analysis in the CFX Manager software (Bio-Rad Laboratories). The optimal annealing temperature was identified using a thermal gradient (50-60°C). Final thermocycling conditions were 2 min at 98°C, 40 cycles of 5 sec at 98°C and 10 sec at the optimal annealing temperature, followed by a melt-curve analysis between 55-95°C in 0.5°C increments. After the gradient qPCR test, each assay was evaluated for specificity against nine other genera using the same PCR conditions (Table 1) prior to further testing using ddPCR. PCR products resulting from successful amplifications were submitted to a commercial facility (Macrogen) for bidirectional Sanger sequencing using the same primers. The resulting sequences were assembled and quality checked using Geneious Prime and matched to reference sequence datasets from NCBI, to test for specificity.

### 2.3 Test conditions-ddPCR

Each ddPCR reaction included 0.5µM of each primer, 1X EvaGreen Supermix (Bio-Rad), and 3µl of DNA template in a total reaction volume of 24 µl. A Bio-Rad AutoDG system was used to automate droplet generation prior to thermal cycling at 95°C for 5 minutes, 39 cycles of 95°C for 30 seconds and the optimized annealing temperature (Table 3) for 1 minute, with a final extension of 4°C for 5 minutes and 90°C for 5 minutes. Droplets were scanned on a QX200 droplet reader (Bio-Rad), where the number of positive and negative droplets per sample were determined by setting a fluorescence threshold, as described below (section 3.1).

### 2.5 Specificity Tests

Tests of analytical specificity were conducted for each primer pair, to ensure amplification only occurred for the target genus and establish the frequency of false positives. Genus-level specificity tests were conducted at the annealing temperature identified during optimization (Table 1) using ten genera (Suppl. Table 1). The genera samples (Suppl. Table 1) used during the specificity tests were extracted using the NucleoSpin® Plant II Kits and were sent off to Macrogen Inc. for sequencing after Sanger sequencing. These steps were done to ensure that assays did not cross-amplify to non-target genera. A BLAST search was then conducted to identify the genera of each non-target sample (Supp. Table 1). Once completed, the target genus, samples of the nine other genera, and a negative control were extracted and tested with the QX200. Each sample was normalized to 1 ng/µl before ddPCR analysis. Five replicate reactions were generated under the ddPCR conditions described above. Test results were analysed for evidence of cross-amplification for other species.

### 2.6 Sensitivity Tests

Sensitivity tests were used to identify the lower limit of detection (LoD) of an assay. A dilution series of 1:10^1^-1:10^5^ was created using gDNA extracted from identified samples, of eDNA extracts from filtered seawater obtained from kelp-free aquaculture tanks at the National Institute of Water and Atmospheric Research (NIWA) Bream Bay to make the diluents. Each sensitivity test was run in triplicate (Agency, 2009; reaserch, 2020; Wissel et al., 2022) at its designated annealing temperature, using the reaction conditions described previously. Thresholds were set directly below the positive droplet cluster for each diluted sample to quantify the concentration (copies/µl)(Figure 1). (Fig. 1; Suppl. Table 2).

**Figure 1.**
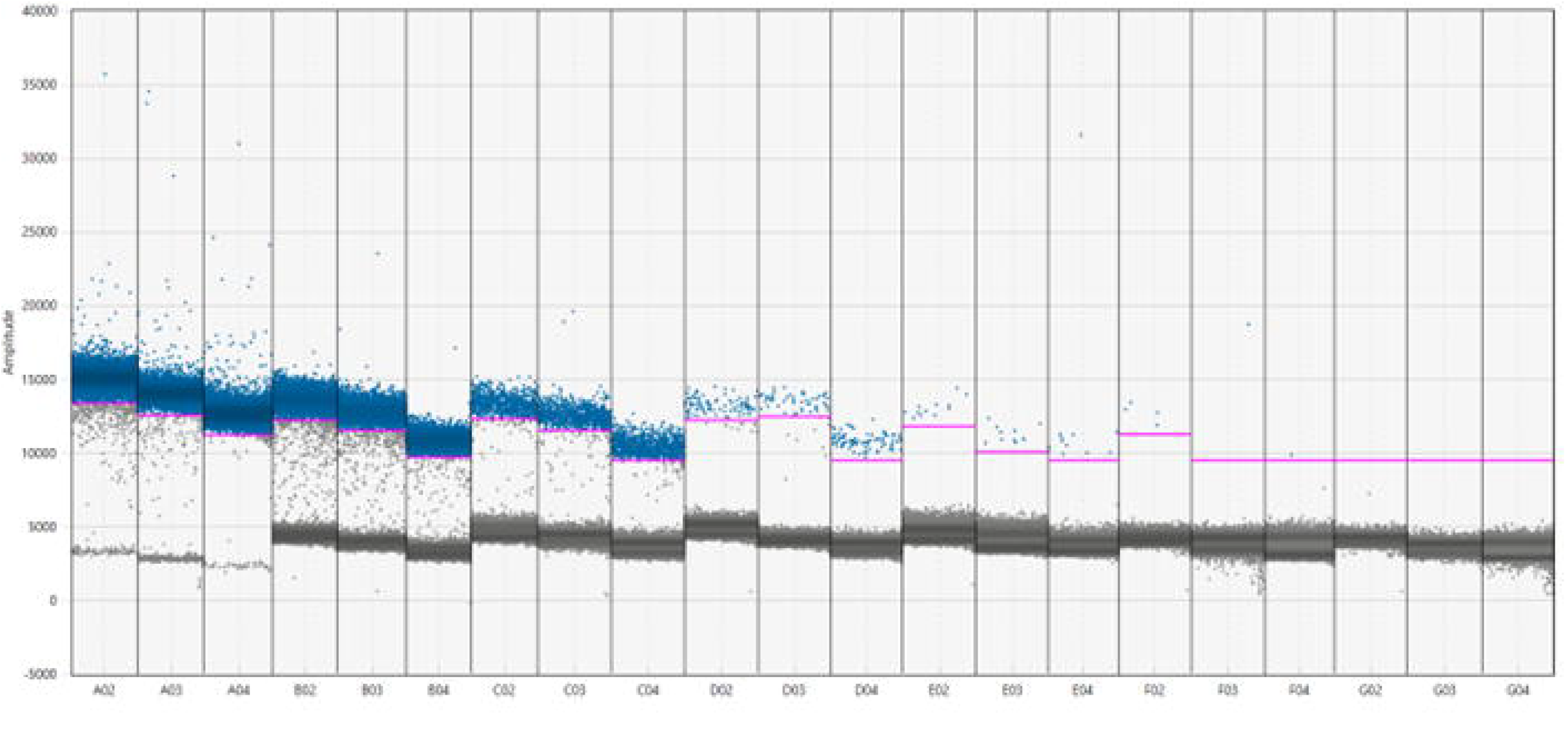
Sensitivity test for M. pyrifera assay. Blue dots indicate the positive droplets being amplified and grey dots are the negative amplitudes during the sensitivity test. The pink line are the individual thresholds for positive amplitudes of each sample. X-axis indicates the sample well and y-axis is the amplitude number

## 3. Results

### 3.1 Assay development

Five ddPCR assays targeting the *cox*3 gene for *Carpophyllum, Cystophora, Ecklonia, Lessonia*, and *Macrocystis* species were successfully designed and validated (Fig. 2; Table 2). The relative fluorescence units (RFU) for both qPCR and ddPCR varied between genera. The RFU values observed during sensitivity testing established the fluorescence threshold for qPCR as 3000 for *Carpophyllum*, 3000 for *Cystophora*, 4000 for *Ecklonia*, 1400 for *Lessonia*, and 4000 for *Macrocystis*. The RFU for positive clusters during ddPCR analysis ranged from 1,000-10,000. Each assay had varying ranges for positive and negative clusters, with some showing droplet ‘rain’. *Carpophyllum* had positive RFU values at 10,000. *Cystophora* had positive RFU values at 23,000. *Ecklonia* positive RFU values at 25,000 *Lessonia* had positive RFU value is at 25,000. *Macrocystis* also shared a positive RFU value 25,000.

**Figure 2.**
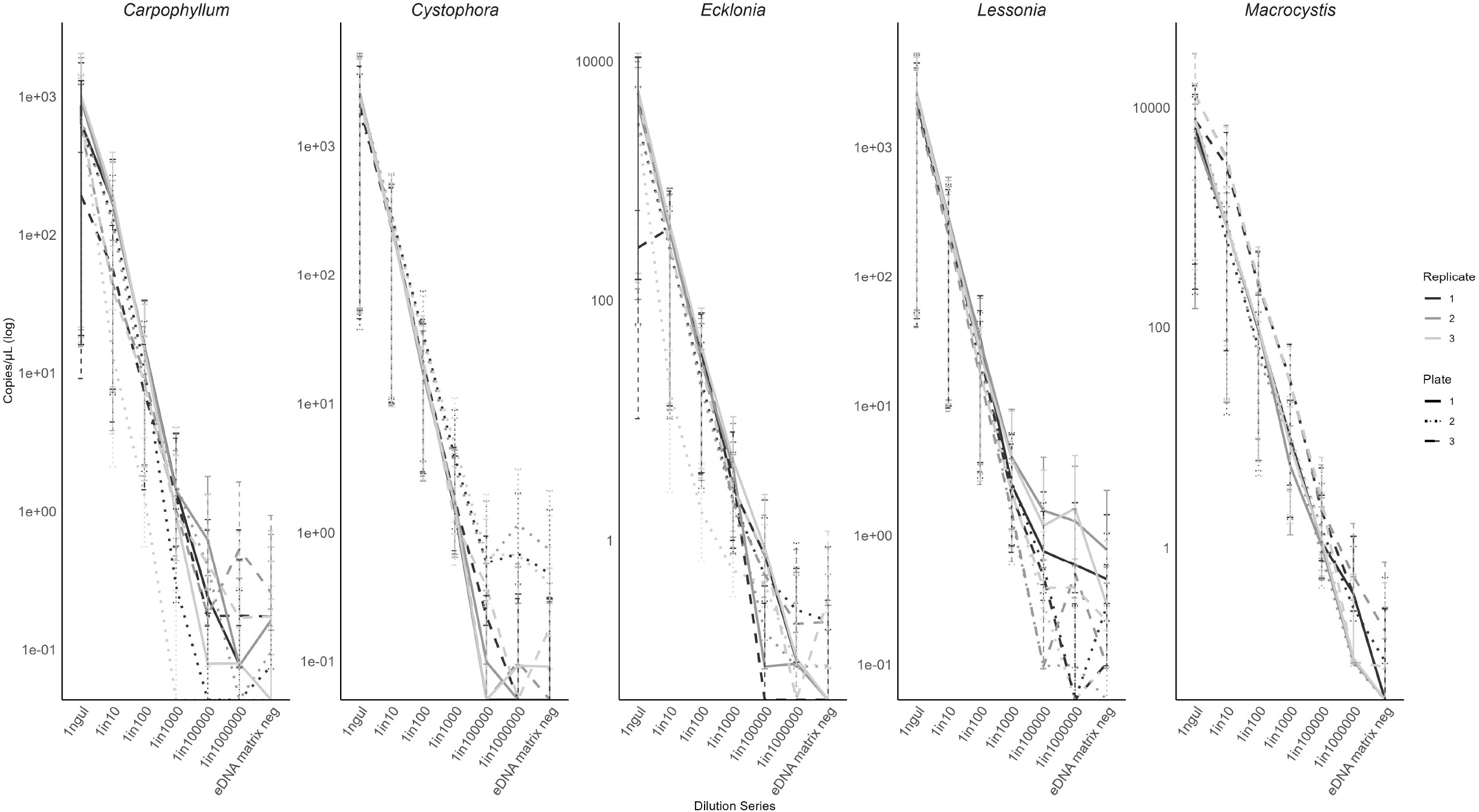
The limit of detection (LoD) for each assay during the sensitivity dilution series. Colours represent the replicate number, 1-3 and line type represents the plate number for that genus, 1-3. Error bars are the minimum and maximum Poisson confidence interval. X-axis represent the dilution and y-axis is the logged copy/µl to estimate the LoD.

### 3.2 Analytical Performance

Each assay was tested with the nine non-target brown algal that had been previously extracted of brown macroalgal genera to validate specificity at the genus level. No cross amplification was observed during the specificity tests, indicating each primer assays is genus-specific. Sensitivity tests were conducted for each of the five assays, to determine the limit of detection (LoD) for each assay. Assays had LoDs of 1.05 copies/µl (SE of 0.0644) for *Carpophyllum*, 2.24 copies/µl (SE of 0.3745) for *Cystophora*, 2.48 copies/µl (SE of 0.3) for *Ecklonia*, 2.38copies/µl (SE of 0.0095) for *Lessonia*, and 1.43 copies/µl (SE of 0.192) for *M. pyrifera* (Fig. 1 & 2; Suppl. Table 2).

The average of the lowest Poisson-derived concentrations of each lowest dilution series with lower 95% confidence interval (CI) that did not overlap with the negative eDNA matrix for *Carpophyllum* was 0.52 cp/µl with 0.01 cp/µl for the eDNA negative control, 1.46 cp/µl for *Cystophora* and 0.19 cp/µl, 1.70 cp/µl for *Ecklonia* and 0.1 cp/µl for the negative control, 1.54 cp/µl for *Lessonia* and 0.32 cp/µl for the negative control, and 0.87 cp/µl for *Macrocystis* and 0.03 cp/µl for the negative control (Suppl. Table 2).

## 4. Discussion

### 4.1 Diagnostic Performance and significance

The five target kelp genera studied here have ecological, historical, and economic importance in NZ (Guiry, 2024; Hurd et al., 2004), as well as other temperate marine regions, such as Australia, South Africa, the West Coast of North America, Chile, Ireland, and Norway, to name a few (Buschmann et al., 2017; Chapman, 1970; González-Roca et al., 2021; Catriona L. Hurd et al., 2023). The assays designed in this study can successfully amplify their targets without cross-amplification and display high sensitivity. The five assays here showed minimal variability between each other, all of which were all in the range of 1-2.5 cp/µl. The thresholds are different between each assay and as pointed out by (Brys et al., 2021) the LoD thresholds vary for other studies. The present study followed diagnostic validation procedures for animal and human health applications (Health, 2021; Hess et al., 2012), which set a strict LoD that produces at least 95% positive replicates (Milbury et al., 2014).

The assays are designed and validated for ddPCR EvaGreen® dsDNA dye oil and no tests were conducted for TaqMan-probe based applications, due to the lower high-throughput costs of EvaGreen-based assays. Therefore, these assays are accessible for labs possessing ddPCR capability and a focus on any of the target genera. The assays may also be applicable for qPCR but would require further validation on that platform. Studies have shown that there is no considerable difference in performance between dye and probe-based amplification (Wang et al., 2022). It has been shown that EvaGreen is just as sensitive in detecting target genes as probe-based systems (Falzone et al., 2020). Thus, we anticipate that similar test performance to that reported here would be obtained if probe-based assays were used instead. However, the use of probe assays may provide additional specificity at the species-level, which would be of value in cases where the discrimination of congeners is necessary.

As stated earlier, the present study used target species from each genus, but did not test for species-specificity. Four of the five tested genera, *Carpophyllum, Cystophora, Ecklonia*, and *Lessonia*, have multiple species members that need to be tested against other species within their genus to establish specificity for the designated species or across the genus. This suggests that some of these assays are only verified as specific within the NZ context (e.g., for *Ecklonia, Lessonia*, and *Cystophora*) and require further testing for verification of species-specificity. For instance, *Lessonia* has 12 accepted species members, with four species present in NZ (Zuccarello & D’Archino, 2024). *Cystophora* is a species rich genus, with members present in NZ and Australia, four of which are present in NZ (Nelson et al., 2023). There are currently five accepted *Carpophyllum* species, all of which are endemic to NZ (Neill & Nelson, 2016; Nelson et al., 2023). Since *Lessonia, Cystophora*, and *Carpophyllum* have more than one species member present in New Zealand, more ddPCR tests would need to be conducted to verify if the assays are species or genus specific. On the other hand, *Ecklonia* consists of seven species members, but *E. radaita* is the only recognized member to occur in NZ (Rothman et al., 2015), subsequently it is likely that this assay is species-specific, but tests verifying this could be done.

### 4.2 Future directions

Interest in using nature-based solutions to mitigate and offset greenhouse gas emissions as carbon credits is increasing (Friess et al., 2022), but the science for mCDR is far behind the industry and demand (Boyd et al., 2024). Using kelp as a C mitigator would require a further expansion of kelp farming (DeAngelo et al., 2023; C. L. Hurd et al., 2023; Wu et al., 2023) with very limited knowledge of the natural role kelp fulfil to mCDR, let alone the consequences that off-shore kelp farming may lead to (Boyd et al., 2024). Since we continue to increase our greenhouse gas emissions and consequently rely on nature-based solutions to offset these emissions, it is therefore necessary to develop methods that are sensitive in quantifying kelp contribution to mCDR.

To understand the presence, distribution, and abundance of kelp in BC and deep-sea environments to verify carbon sequestration, it is essential for eDNA assays to be both sensitive and precise in their detection and quantification, respectively. Such assays can aid in creating ethical regulations and policy making regarding kelp in a BC industry. Furthermore, creating a more robust data inventory of what kelp species detritus may be settling in marine C sinks, could provide evidence of the variations and considerations concerning kelp and mCDR. Since the genera-specific assays developed during this study are designed ddPCR, it is probably to say the assays are sensitive to detect target kelp genera for in-situ samples. Although no quantification was attempted in this study, future studies can quantify these assays, thus being able to estimate how much kelp detritus and DNA settle in ocean C sinks. Furthermore, using ddPCR assays developed in this study is a new step to detect specific kelp genera that could provide valuable insight into their past and present contributions to mCDR, as well as aid in predicting their future roles as climate change progresses.

These requirements are also central for other studies and management efforts focusing on kelp, as well. Prior to this study, there was no scientific literature on targeted eDNA assays specific to kelp species or highly sensitive methods to quantify kelp DNA abundance. The present study has developed and validated five genera-specific kelp ddPCR assays capable of Although these assays were designed for BC studies, they may be promising for other studies focusing on any of the five genera due to their potential sensitivity and specificity, however further diagnostic tests would need to be applied to other species members in each genus for validation. For instance, kelp are facing numerous threats and environmental pressures leading to population declines and changes in their distributional ranges and abundance due to climate change, global temperature increases, and anthropogenic activities (Becheler et al., 2022; Blain et al., 2021; Brodie et al., 2014; Bunting et al., 2024; Coleman et al., 2022; Mason & Blain, 2024; McPherson et al., 2021; Michaud et al., 2022; Smale, 2020; Smale et al., 2013). Therefore, it would be highly beneficial to monitor changes in kelp species abundance, distribution changes, and potential for detritus transportation. The assays designed and validated in the present study may be applied to several avenues of scientific research, monitoring programs, and aquaculture operations that have an interest in the target kelp this study is based on.

The results from the assay validation tests indicate that the assays have the capability to detect kelp DNA at different concentration levels. However, further studies are necessary for the objective of measuring each assay’s ability to quantify target genera. This study did not quantify DNA concentration compared with kelp biomass; however, quantification will be assessed in a different study by the authors.

## 5. Conclusion

Using eDNA methodologies for ecological monitoring and mCDR studies is a valuable, non-invasive, and economically valuable tool. Designing a genus-specific eDNA assay will provide further insights into cryptic environments by determining what species are present, and estimations of how much kelp-specific DNA is in a sample. Assays such as these can be especially useful for understanding and quantifying the contribution that kelp fulfil in mCDR. Conservative thresholds were used in all the assays to provide for minimum estimates of sensitivity. Therefore, the results likely underestimate the sensitivity levels of all five primer assays. Designing genera-specific primer eDNA assays will allow for ecological monitoring of these species on regional and global levels. Along with eDNA ecological monitoring, assays such as those advanced in the present study may be utilized for research that examines the presence and quantification of kelp detritus in marine carbon sinks. Utilizing these assays can assist in identifying kelp transport mechanisms and sequestration pathways once dislodged from their rocky reef coastal habitat.

## Abbreviations

(CO_2_: Carbon dioxide
(mCDR: marine carbon dioxide removal
(C: Carbon
(BC: Blue Carbon
(eDNA: Environmental DNA
(PCR: Polymerase Chain Reaction
(qPCR: real-time PCR
(ddPCR: droplet digital PCR
(*cox*3: cytochrome c oxidase subunit III protein coding gene
(LoD: Limit of detection
(CI: confidence interval
(RFU: relative fluorescence units

