## Supplemental Table for "Development and validation of five native Aotearoa New Zealand genera-specific large brown macroalgal eDNA ddPCR assays"

Supp. Table 1. Brown macroalgal species used for assay design and validation. Columns indicate Identifier number was given to each sample for identification by the authors for this project. GPS coordinates indicate where the samples were collected from the Wellington region.

| Genus | Species | GPS coordinates (latitude, longitude) | Identifier Number | Collection Site | Date collected |
| --- | --- | --- | --- | --- | --- |
| *Carpophyllum* | *maschalocarpum* | -41.346447,  174.791202 | NZBC038 | Princess Bay, Wellington | 22-Dec-22 |
| *Cystophora* | *torulosa* | -41.297700,  174.820211 | NZBC034 | Shelly Bay, Wellington | 21-Dec-22 |
| *Ecklonia* | *radiata* | -41.332545, 174.830077 | NZBC018 | Breaker Bay, Wellington | 10-Jan-23 |
| *Lessonia* | *variegata* | -41.342153  174.791290 | NZBC036 | Waitaha Cove, Wellington | 4-Jan-2023 |
| *Macrocystis* | *pyrifera* | -41.292191, 174.833078 | NZBC035 | Mahanga Bay, Wellington | 22-Nov-21-Dec-22 |
| *Undaria* | *pinnatifida* | -41.342354, 174.809399 | NZBC003 | Moa Point, Wellington | 01-Dec-2022 |
| *Marginariella* | *boryana* | -41.613513, 175.291853 | NZBC001 | Cape Palliser | 03-Nov-22 |
| *Marginariella* | *urviliana* | -41.613513, 175.291853 | NZBC002 | Cape Palliser | 03-Nov-22 |
| *Tinocladia* | *novae zealandia* | -41.342354, 174.809399 | NZBC011 | Moa Point, Wellington | 01-Dec-2022 |
| *Durvillaea* | *antarctica* | -41.342655, 174.809820 | NZBC039 | Moa Point, Wellington | 01-Dec-2022 |
| *Lobophora* | *sp.* | -41.342354, 174.809399 | NZBC012 | Moa Point, Wellington | 01-Dec-2022 |
| *Dictyota* | *kuntii* | -41.342354, 174.809399 | NZBC013 | Moa Point, Wellington | 01-Dec-2022 |

Supp. Table 2. This table are the average values of copies/µl, Poisson Confidence max and min for each dilution seires of the five assays.

| **Target Species** | **Dilution** | Concentration (copies/µL) (avg) | Poisson Confidence max (avg) | Poisson Confidence min (avg) |
| --- | --- | --- | --- | --- |
| *Carpophyllum maschalocarpum* | 1 ng/µl | 694.05 | 658.15 | 626.7 |
| *Carpophyllum maschalocarpum* | 1:10 | 113.56 | 100.48 | 90.09 |
| *Carpophyllum maschalocarpum* | 1:100 | 10.58 | 11.02 | 7.77 |
| *Carpophyllum maschalocarpum* | 1:1000 | 1.05 | 1.53 | 0.52 |
| *Carpophyllum maschalocarpum* | 1:10000 | 0.22 | 0.58 | 0.07 |
| *Carpophyllum maschalocarpum* | 1:100000 | 0.11 | 0.43 | 0.03 |
| *Carpophyllum maschalocarpum* | eDNA neg | 0.1 | 0.42 | 0.02 |
| *Cystophora retroflexa* | 1 ng/µl | 2146.54 | 2200 | 2108 |
| *Cystophora retroflexa* | 1:10 | 237.68 | 247.45 | 228.55 |
| *Cystophora retroflexa* | 1:100 | 22.39 | 25.1 | 19.57 |
| *Cystophora retroflexa* | 1:1000 | 2.24 | 3.15 | 1.46 |
| *Cystophora retroflexa* | 1:10000 | 0.34 | 0.68 | 0.09 |
| *Cystophora retroflexa* | 1:100000 | 0.29 | 0.68 | 0.12 |
| *Cystophora retroflexa* | eDNA neg | 0.19 | 0.57 | 0.07 |
| *Ecklonia radiata* | 1 ng/µl | 3815.02 | 3755.34 | 3557.66 |
| *Ecklonia radiata* | 1:10 | 318.47 | 302.26 | 282.15 |
| *Ecklonia radiata* | 1:100 | 26.7 | 27.83 | 22.3 |
| *Ecklonia radiata* | 1:1000 | 2.59 | 3.48 | 1.7 |
| *Ecklonia radiata* | 1:10000 | 0.43 | 0.89 | 0.17 |
| *Ecklonia radiata* | 1:100000 | 0.13 | 0.43 | 0.01 |
| *Ecklonia radiata* | eDNA neg | 0.1 | 0.45 | 0.02 |
| *Lessonia variegata* | 1 ng/µl | 2221.93 | 2243.47 | 2152.2 |
| *Lessonia variegata* | 1:10 | 221.74 | 234.41 | 216.1 |
| *Lessonia variegata* | 1:100 | 22.49 | 24.73 | 19.26 |
| *Lessonia variegata* | 1:1000 | 2.47 | 3.27 | 1.52 |
| *Lessonia variegata* | 1:10000 | 0.7 | 1.09 | 0.25 |
| *Lessonia variegata* | 1:100000 | 0.48 | 0.94 | 0.22 |
| *Lessonia variegata* | eDNA neg | 0.32 | 0.65 | 0.07 |
| *Macrocystis pyrifera* | 1 ng/µl | 6282.74 | 8717.66 | 9654.46 |
| *Macrocystis pyrifera* | 1:10 | 1422.17 | 1500.19 | 1432.98 |
| *Macrocystis pyrifera* | 1:100 | 125.26 | 132.76 | 120.04 |
| *Macrocystis pyrifera* | 1:1000 | 14.76 | 17.25 | 13.01 |
| *Macrocystis pyrifera* | 1:10000 | 1.43 | 2.22 | 0.87 |
| *Macrocystis pyrifera* | 1:100000 | 0.19 | 0.59 | 0.05 |
| *Macrocystis pyrifera* | eDNA neg | 0.03 | 0.31 | 0 |
